# A robust workflow for 3D imaging of human mitochondria using cryo-electron tomography

**DOI:** 10.64898/2026.04.16.717704

**Authors:** Akhil Gargey Iragavarapu, Oleg Artemchuk, Daija Bobe, Anna C. Ratliff, Evgeny Pavlov, Halil Aydin

**Affiliations:** Department of Molecular Pathobiology, College of Dentistry, New York University, New York, NY, USA; New York Structural Biology Center, New York, NY, USA; Midwest Center for Cryo-Electron Tomography, Madison, WI, USA; Pain Research Center, New York University, New York, NY, USA; Department of Biochemistry and Molecular Pharmacology, Grossman School of Medicine, New York University, New York, NY, USA

## Abstract

Mitochondria are dynamic signaling organelles that transduce metabolic and biochemical cues to facilitate cellular adaptation. Their complex structure and dynamics are essential for integrating metabolic pathways, responding to stressors, and communicating inter- and intra-cellular signals. While optimal mitochondrial activity is frequently linked to cellular and organismal health—influencing processes ranging from metabolism and regulated cell death to differentiation and growth—the mechanistic links between mitochondrial dysfunction and cellular defects leading to human disease remain incompletely understood. Understanding how mitochondrial shape and function are linked is crucial for deciphering the regulatory mechanisms of cell survival and fate. Here, we present a molecular resolution cryo-electron tomography (cryo-ET) imaging and image analysis platform to investigate the structure of isolated human mitochondria under different conditions. We describe optimized protocols for isolating mitochondria from human cells, vitrifying these samples with high-pressure freezing (HPF) using the waffle method, cryo-focused ion beam (cryo-FIB) milling to generate thin sections (lamellae), and imaging with cryo-transmission electron microscopy (cryo-TEM). This is complemented by a robust downstream processing pipeline for tilt-series alignment, tomogram reconstruction, and three-dimensional (3D) segmentation of tomograms using the latest state-of-the-art algorithms. With some variations, this versatile workflow is adaptable to other subcellular compartments for structural studies in isolation or within intact cells. Furthermore, our protocols provide a critical foundation for investigating the *in-situ* structure of protein machineries that govern key cellular processes.

## 1. Introduction

Eukaryotic cells contain distinct membrane-bound and membraneless organelles that enable multiple, often incompatible, chemical reactions to occur simultaneously within the same cell (Vosseberg et al., 2024). Each organelle consists of hundreds of different macromolecules, such as proteins, lipids, nucleic acids, and metabolites, which form functional networks that underlie many cellular processes, including gene expression, metabolism, signaling, differentiation, and growth (Carlton et al., 2020; Van Bergeijk et al., 2016). While the spatial and temporal organization of organelles and their macromolecules impact the behavior and function of eukaryotic cells, understanding how these subcellular compartments perform specific tasks still remains incompletely understood.

Mitochondria are major signaling organelles that regulate metabolic, stress response, survival, and growth pathways within the cell (Shen et al., 2022). Originating from an alphaproteobacterial lineage that became a permanent endosymbiont in an archaeal host, primitive mitochondria relinquished almost all of their genes within the ancestral eukaryotic cell (Roger et al., 2017; Youle, 2019). Although human mitochondria are comprised of ∼1200 proteins, the small mitochondrial genome only encodes 13 proteins of mitochondrial respiratory complexes, and approximately 99% of mitochondrial proteins reside in nuclear-encoded genes that are translated in the cytosol and imported into the mitochondrion (Youle, 2019). The mitochondrial protein machines perform a wide variety of functions, including ATP production via oxidative phosphorylation (OXPHOS), metabolism of amino acids, lipids, and nucleotides, regulation of cellular calcium homeostasis, control of reactive oxygen species (ROS), and programmed cell death (Bennett et al., 2022; Kamerkar et al., 2025; Manicki et al., 2022; Pfanner et al., 2019; Spinelli and Haigis, 2018). The functional diversity of mitochondrial protein networks is also reflected in mitochondrial ultrastructure (Tábara et al., 2025). Mitochondria are partitioned into multiple subcompartments by an inner and an outer membrane (Daumke and Van Der Laan, 2025; Muthukumar and Weissman, 2025). The inner membrane folds inwards to form invaginations called cristae, which are highly dynamic compartments that increase the surface area of the membrane to accommodate many copies of the proteins and enzymes involved in the OXPHOS system (Daumke and Van Der Laan, 2025; Dietrich et al., 2024; Pfanner et al., 2019; Zheng et al., 2024). Additionally, mitochondria are highly dynamic organelles and adopt divergent morphologies in different cell types, phases of the cell cycle, and stress stimuli (Archer, 2013). These dynamic changes in mitochondrial morphology, controlled by several mitochondria-shaping proteins, occur in response to both internal and environmental signals, and tune the function of the organelle according to physiological demands (Cao et al., 2017; Kalia et al., 2018; Liu et al., 2024; Von Der Malsburg et al., 2023; Yamada et al., 2025).

Defects in mitochondrial processes are often accompanied by the progressive deterioration of cell structure and function, leading to the development of numerous human diseases, including neurodegenerative disorders, metabolic diseases, cancer, and age-related illnesses (Jung et al., 2025; Russell et al., 2020; Thatavarthy et al., 2025; Wu et al., 2023). Mitochondrial dysfunction is characterized by a systemic collapse of bioenergetic homeostasis, primarily driven by the uncoupling of the electron transport chain (ETC) and the dissipation of the mitochondrial membrane potential (Glover et al., 2024). This disruption leads to a critical decline in the ATP/ADP ratio, impairing essential cellular processes such as protein synthesis and the maintenance of ion gradients (Glover et al., 2024). When mitochondria experience inordinate stress and lose function, they also generate metabolites and reactive oxygen species (ROS) as by-products that activate stress signals and can be toxic to the cell (Schieber and Chandel, 2014). Loss of mitochondrial inner membrane integrity triggers the formation of the mitochondrial permeability transition pore (mPTP), leading to mitochondrial swelling and the efflux of ROS, Ca^2+^, mitochondrial DNA (mtDNA), and pro-apoptotic proteins like cytochrome *c* into the cytosol (Bernardi et al., 2023; Bonora et al., 2022; Neginskaya et al., 2022, 2019). Additionally, the failure of mitophagy results in the accumulation of dysfunctional mitochondria, exacerbating cellular proteotoxicity (Picca et al., 2023; Uoselis et al., 2023). Together, these biochemical perturbations become particularly catastrophic in post-mitotic tissues with high energy demand, such as neurons and cardiomyocytes, manifesting as the diverse clinical phenotypes of mitochondria-associated diseases (Archer, 2013; Russell et al., 2020).

Quantitative explanations of mitochondrial mechanisms should bridge length scales from organelles and their internal structures, like cristae (microns), to nearest-neighbor protein interactions (nanometers) and provide synergistic evidence for the mechanisms that modulate mitochondrial output (Barad et al., 2023; Beck et al., 2024; Nogales and Mahamid, 2024). Recent advances in cryo-electron tomography (cryo-ET) imaging technologies enable the visualization of intact organelles and the structures of even the most intricate machines at molecular-resolution detail (Anton et al., 2025; Mahamid et al., 2016; Singh et al., 2024; Von Appen et al., 2015; Waltz et al., 2025). Cryo-ET requires samples to be rapidly frozen before they are imaged at cryogenic temperatures (Kaplan et al., 2021; Klykov et al., 2022). Vitrification of biological samples can be achieved using different techniques (De Winter et al., 2021; Kelley et al., 2022; Klykov et al., 2022). While a semi-automated plunge freezer, such as the Vitrobot Mark IV System (Thermo Fisher Scientific) or EM GP2 Automatic Plunge Freezer (Leica Microsystems), can freeze samples up to ∼10 µm thick, a high pressure freezing (HPF) system can allow for the vitrification of samples up to ∼200 µm in thickness (Kaplan et al., 2021). Vitrified samples are then subjected to cryo-correlative light and electron microscopy (cryo-CLEM) or integrative Fluorescence Light Microscope (iFLM) to identify the localization of fluorescently labeled organelles or proteins (Yang et al., 2026). Subsequently, cryo-focused ion beam milling (cryo-FIB) is used to mill these targets into ∼150 to 250 nm thin sections (lamellae) (Thermo Fisher Scientific Aquilos 2), enabling electron penetration and a higher signal-to-noise ratio (SNR) of the data (Berger et al., 2023; Kuba et al., 2021). Cryo-FIB milled thin sections can be imaged using a cryo-transmission electron microscope (cryo-TEM) operated at 300 KeV (Thermo Fisher Scientific Krios) and a direct electron detector (Thermo Fisher Scientific Falcon 4i or Gatan K3) equipped with an imaging filter (Thermo Fisher Scientific Falcon Selectris or Gatan BioContinuum). Current automated, high-voltage cryo-TEMs direct electron detectors allow for the acquisition of high-throughput and high-quality images (micrographs) of the biological specimen from different angles (from -60° to +60° with 1°–3° tilt increments), and enhance the SNR of the tilt images, while grouped dose-symmetric data acquisition strategy provide ideal dose distribution for obtaining high-resolution reconstructions (Hagen et al., 2017; Mastronarde, 2005; Turoňová et al., 2020). The resulting tilt series are then motion corrected, aligned to correct for spatial displacements during data acquisition, and used for calculating the reconstruction of tomograms, which provide three-dimensional (3D) representation of the specimen (Burt et al., 2024; Chen et al., 2019; Mastronarde and Held, 2017; Peck et al., 2025a). At this stage, segmentations of specific features and subtomogram averaging (STA) of repetitive objects can be performed to determine their 3D structures and properly interpret high-resolution structural signals from images (Chen et al., 2019; Wan and Briggs, 2016).

In this protocol, we present a general workflow to elucidate the nanometer-scale organization and dynamics of isolated human mitochondria using cryo-ET. We combine detailed descriptions of sample preparation and imaging steps with the most recent computational tomogram reconstruction and segmentation methods to provide a comprehensive guide for cryo-ET analysis. This robust pipeline should be adaptable to other subcellular compartments and *in situ* investigations of intact cells and can be used for determining the molecular architecture of cellular machines.

## 2. Materials

### 2.1 Common disposables

1. Pipettors and tips 10 μL, 200 μL, 1,000 μL (Rainin, catalog no. 30389267, 30389186, and 30389260)
2. Conical Sterile Polypropylene Centrifuge Tubes (Thermo Fisher Scientific, catalog no. 339652)
3. Kimwipe tissues (Kimtech, catalog no. 34155)
4. Filter papers (Whatman, catalog no. 1004090)
5. T75 adherent cell culture flasks (Thermo Fisher Scientific, catalog no. 156800)
6. Abrasive sandpaper with grits 1,200, 7,000, 15,000, or similar (ADVcer, catalog no. ADV Sandpaper Sheets S01 5 10BUGY)

### 2.2 Cells, reagents, and chemicals

1. Human HAP1 cells (Horizon, catalog no. C631)
2. Dulbecco’s Modified Eagle Medium (DMEM) (Sigma Aldrich, catalog no. D6429)
3. Fetal bovine serum (FBS) (Sigma Aldrich, catalog no. F2442)
4. Penicillin-streptomycin (Sigma Aldrich, catalog no. P4333)
5. Trypsin-EDTA (Thermo Fisher Scientific, catalog no. 25200072)
6. MitoTracker™ Dyes for Mitochondria Labeling (Thermo Fisher Scientific, catalog no. M7514)
7. HBSS solution (HBSS GIBCO, catalog no. 14025092)
8. Fatty acid free bovine serum albumin (Sigma Aldrich, catalog no. A6003)
9. Mannitol (Sigma Aldrich, catalog no. PHR1007)
10. EGTA (Sigma Aldrich, catalog no. E3889)
11. Sucrose (Sigma Aldrich, catalog no. S0389)
12. HEPES (Sigma Aldrich, catalog no. H3375)
13. 1-Hexadecene (Millipore Sigma, catalog no. 822064)
14. Metal-polish WENOL (Ted Pella, catalog no. 892)

### 2.3 Equipment

1. GKH GT Motor Control (Glas-Col, catalog no. 099C K44)
2. Gridboxes (SubAngstrom, catalog no. GBV01)
3. Aluminum planchettes Type B 0.3 mm recess (Ted Pella, catalog no. 39201)
4. Quantifoil R2/4 Au 200-mesh cryo-EM grids (SPI Supplies, catalog no. 4220C-CF)
5. SEM pin stub (Ted Pella, catalog no. 16261)
6. HPF holder (Technotrade International, catalog no. 290-1)
7. C-clip ring (Thermo Fisher Scientific, catalog no. 1036173)
8. Cryo-FIB AutoGrid (Thermo Fisher Scientific, catalog no. 1205101)
9. Clipping station and clipping tools (Thermo Fisher Scientific, catalog no. 1000068)
10. Centrifuge (Thermo Fisher Scientific, catalog no. 75004240)
11. Tweezers (Ted Pella, catalog no. 5220)
12. AutoGrid tweezers (Ted Pella, catalog no. 47000-600)
13. Personal protective equipment for work with liquid nitrogen (gloves, eye protection, oxygen monitors)
14. Grid transfer dewar (Thomas Scientific, catalog no. 5028M40)
15. Industrial grade dry nitrogen tank
16. Industrial grade liquid nitrogen tank
17. Liquid Nitrogen dewar with 4 L capacity
18. Liquid Nitrogen dewar or puck system for grid storage (SubAngstrom, catalog no. GSS02-E)
19. CCU-010 HV high vacuum coater (Safematic, catalog no. 100001)
20. CT-010 carbon fiber evaporation head module (Safematic, catalog no. 100003)
21. Plasma Cleaner (Gatan Inc., Gatan Solarus II Model 955)
22. Wohlwend Compact 01 HPF or comparable (M. Wohlwend GmbH)
23. Zeiss LSM 900 Airyscan confocal microscope with a Linkam cryo stage
24. Cryo-FIB/SEM dual beam microscope, including relevant grid shuttles (Thermo Fisher Scientific, Aquilos 2)
25. Anti-contamination cryo-shield/cryo-shutter system (Delmic CERES Ice Shield, catalog no. DM-2707-999-0003-1)
26. Titan Krios G4 TEM (Thermo Fisher Scientific) with a cold Field Emission Gun (E-CFEG), a SelectrisX imaging filter operated with a 10-eV slit width, and a Falcon 4i direct electron detector

### 2.4 Software

1. Software for milling automation. AutoTEM Cryo version 2.2 (Kuba et al., 2021).
2. Microscope operating platform (Thermo Fisher Scientific, Aquilos 2) xT version 20.1.1.
3. Automated cryo-ET data acquisition (Thermo Fisher Scientific, Titan Krios G4 TEM), SerialEM version 4.1.0 (Mastronarde, 2005).
4. Beam-induced motion correction on raw movie frames, MotionCor2 version 1.6.4 (Zheng et al., 2017).
5. Contrast transfer function (CTF) parameters, CTFFIND4 (Rohou and Grigorieff, 2015).
6. Automated tomographic reconstruction, IMOD version 5.1 (Mastronarde and Held, 2017).
7. Align frames from dose-fractionated images, AlignFrames version 4.12.62 within IMOD (Mastronarde and Held, 2017).
8. Edit image file header metadata, Alterheader version 4.12.62 within IMOD (Mastronarde and Held, 2017).
9. Make movies from image outputs, FFMPEG version 5.1.2.
10. Motion correction, CTF correction, Tilt-series alignment, and Tomogram reconstruction, AreTomo3 version 2.2.2 (Peck et al., 2025a).
11. Tomogram denoising and missing-wedge correction, Isonet2 version 2.0.0 (Liu et al., 2025)
12. Automated tomogram segmentation, MemBrain v2 (Lamm et al., 2025).
13. Data visualization, IMOD version 4.11.25 (Kremer et al., 1996) and ChimeraX version 1.11.1 (Meng et al., 2023).

## 3. Methods

### 3.1 Cell culture and mitochondria isolation

Human HAP1 cells are grown under standard adherent mammalian cell culturing conditions (37 °C with 5% CO_2_) for mitochondrial isolation. Before isolation, mitochondria are fluorescently labeled in intact cells with MitoTracker green to identify the units of organelle during downstream sample preparation and imaging. After staining, cells are harvested and washed in ice-cold isolation buffer to preserve mitochondrial integrity. Mitochondria are then isolated by gentle mechanical homogenization followed by differential centrifugation to separate the organelle from intact cells, nuclei, and other cellular debris. A sucrose-based buffer is used for final wash and resuspension to further improve sample purity while maintaining osmotic stability. The purified mitochondrial pellet is kept on ice and used for grid preparation within a few hours for optimal sample quality.

1. Grow HAP1 cells in Dulbecco’s Modified Eagle Medium (DMEM) supplemented with 10% fetal bovine serum (FBS) and 1% penicillin-streptomycin (or antibiotic-antimycotic) in a humidified incubator at 37°C with 5% CO₂.
2. Maintain cells as adherent cultures in T75 flasks (75 cm² growth surface area, typically with 15–20 mL of culture medium per flask).
3. Passage at 70–90% confluency using 0.05% trypsin-EDTA to detach them, typically every 2–4 days depending on growth rate.
4. Fluorescently label mitochondria within the intact cells using MitoTracker green.
5. Remove the culture medium and wash the attached cells twice with standard phosphate-buffered saline (PBS).
6. Cover the cells with 10 mL of HBSS solution containing 100 nM MitoTracker green and incubate in the dark for 20 minutes at 37°C with 5% CO_2_.
7. After staining, harvest the cells at approximately 80–90% confluency from multiple T75 flasks by trypsinization or gentle detachment and centrifuge at 400x *g* for 5 minutes to obtain the cell pellet.
8. Rinse the cell pellet three times in ice-cold isolation buffer containing 5 mM HEPES (pH 7.2), 225 mM mannitol, 75 mM sucrose, and 1 mM ethylene glycol tetraacetic acid (EGTA) supplemented with 1 mg/mL fatty acid-free bovine serum albumin.
9. Resuspend the cell pellet in ice-cold isolation buffer and homogenize on ice in a 5 mL Teflon-glass homogenizer using 40 gentle strokes of the pestle rotating at 40 rpm using a Glas-Col GKH GT Motor Control (*see Note 1*).
10. Spin the homogenate at 1000x *g* for 10 min at 4°C to remove intact cells and debris.
11. Transfer the supernatant to a new tube and spin the cells at 10,000 x *g* for 10 minutes at 4°C.
12. Discard the supernatant and resuspend the mitochondrial pellet in 500 μL of ice-cold isolation buffer or a sucrose buffer containing 5 mM HEPES (pH 7.2), 250 mM sucrose, and 100 µM EGTA.
13. Centrifuge again at 10,000x *g* for 10 min at 4°C to further purify the mitochondria.
14. After the final centrifugation, discard the supernatant, and cover the purified mitochondrial pellet with 100 μL of sucrose buffer.
15. Keep the samples on ice in a pellet form until vitrification within 2 to 3 hours.

### 3.2 Cryo-electron tomography sample preparation

Cryo-electron tomography (cryo-ET) sample preparation begins with reinforcing EM grids and preparing planchettes to support reliable waffle formation during high-pressure freezing (HPF). The grids are coated with an additional layer of carbon to improve rigidity, while the planchettes are polished and treated with 1-hexadecene to promote clean separation after vitrification. The sample is loaded between coated planchettes on the grid to create the waffle configuration for freezing (*see Note 2*). The waffle assembly is then vitrified in a high-pressure freezer, which preserves native ultrastructure by preventing ice crystal formation. After freezing, the waffle assembly is carefully recovered under liquid nitrogen, the planchettes are removed, and the vitrified specimens are clipped into AutoGrids with defined orientation for downstream cryo-FIB/SEM processing (*see Note 3*).

1. To reinforce grids with an extra carbon layer, place the carbon side of the grid facing up on a clean glass slide (*see Note 4*).
2. Transfer the glass slide into a carbon evaporator and deposit ∼20 nm of carbon onto the carbon side of the R2/4 Au 200-mesh EM grids (*see Note 5*).
3. Sand the flat side of the Type B planchettes sequentially using 1,200, 7,000, and 15,000 grit sandpaper to remove manufacturing rings.
4. Polish the flat surface using UNIPOL metal polish with circular motions on a filter paper and remove excess polish using a Kimwipe.
5. Perform a final rub on clean filter paper to ensure the surface is residue-free.
6. Place the polished planchettes flat side up on filter paper.
7. Add 4–6 μL of 1-Hexadecene onto the surface of each planchette, spread the droplet across the surface using the pipette tip, and then apply a second droplet to ensure full coating.
8. Allow the coated planchettes to sit for at least 15 minutes before waffle assembly.
9. Plasma clean the carbon-coated grids with the carbon side down for 30 s using 80% O₂ and 20% H₂ at 50 W radio frequency (RF).
10. Blot away the excess 1-Hexadecene from the coated planchettes using a filter paper.
11. Place one planchette flat side up on the HPF holder tip and position the carbon side of the grid down on the planchette.
12. Add 5 µL of the sample onto the grid.
13. Place a second planchette flat side down on top of the assembly, close the HPF tip, and tighten the assembly using tweezers (*see Note 6*).
14. Load the fully assembled sample into the chamber of a high-pressure freezer.
15. Vitrify the sample at a typical operating pressure of ∼2,100 bar.
16. After vitrification, transfer the HPF holder into the liquid nitrogen chamber.
17. Loosen the holder tip and take the waffle assembly out of the holder.
18. Gently remove the planchettes from the grid (*see Note 7*).
19. Prepare the grid clipping setup used for semi-automated cryo-FIB/SEM workflow and position a stereoscope above the clipping station (*see Note 8*).
20. Label the AutoGrid notch with a marker to aid grid bar alignment during clipping and to help orient lamellae relative to the TEM stage tilt axis during grid loading.
21. Place the grid into the AutoGrid holder under the stereoscope.
22. Rotate the grid using tweezers until the grid squares are aligned with the AutoGrid notch.
23. Secure the grid with the C-clip ring (*see Note 9*).

### 3.3 Cryo-fluorescence screening of grids

Cryo-fluorescence microscopy analysis was performed to identify regions containing fluorescently labelled mitochondria on vitrified grids before cryo-FIB milling and cryo-ET imaging. Frozen grids were imaged using a Zeiss LSM 900 Airyscan confocal microscope equipped with a Linkam cryo stage.

1. Connect the Linkam cryo stage to power.
2. Begin cooling by pouring liquid nitrogen into the reservoir, then press the control button to allow nitrogen to flow from the reservoir into the sample chamber.
3. Ensure that the sample cassette and cassette holder are fully cooled before loading grids.
4. Open the self-aligning magnetic sample cassette within the cassette holder and load up to three unclipped grids.
5. Close the sample cassette and transfer it to the cryo stage using the magnetic transfer tool.
6. Attach the Linkam cryo stage to the microscope.
7. Adjust the Z position and navigate to a grid using the live reflection channel in the Zeiss acquisition software.
8. Switch to the Enhanced Green Fluorescence Protein (EGFP) channel and adjust the focus.
9. Assign a name to the grid and select a folder for automatic file saving.
10. Acquire an image of the grid with both reflection and EGFP channels selected at no magnification, 5x, and 100x magnifications (Figure 1A).
11. Update the file name for each grid before acquiring additional images.

**Figure 1.**
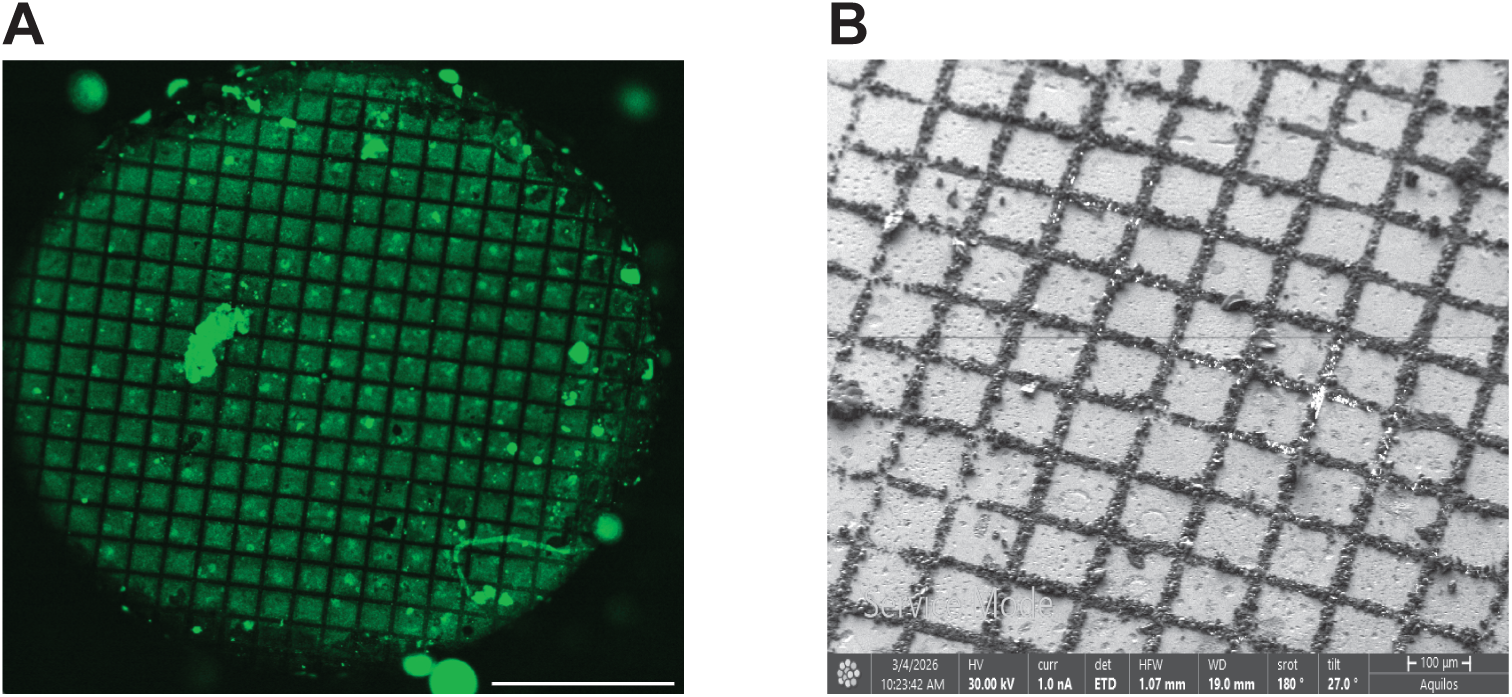
Screening frozen grids prior to cryo-FIB milling. **(A)** Cryo-fluorescence microscopy image of the frozen hydrated grid used to identify the location of the fluorescently labeled specimen. Scale bar, 500 μm. **(B)** A representative image of the ion beam overview after deposition of an organometallic platinum protective layer by gas injection system (GIS) prior to milling.

### 3.4 *Cryo-*focused ion beam milling

Cryo-focused ion beam (Cryo-FIB) milling is used for thinning the fluorescently active regions of the vitrified specimens into electron-transparent lamellae suitable for cryo-electron tomography. The workflow entails system preparation, grid mapping, deposition of a protective organometallic platinum layer onto the sample surface to stabilize it, and ion-beam milling. Within each frozen grid, candidate sites for lamella preparation are identified, inspected, and precut to explore suitable regions and determine the geometry needed for downstream thinning. After remapping the grid, lamella positions are refined and imported into AutoTEM, where guided milling is used to perform bulk removal, angle adjustment, and progressive thinning. Additional notch milling is introduced to provide lamella stress relief and mechanical stability for subsequent handling and imaging steps. Lastly, lamella shaping and optional polishing steps are performed to obtain lamellae of appropriate thickness and quality for cryo-ET image acquisition.

1. Before loading the specimen, purge the GIS lines while the FIB/SEM system is cooling down (*see Note 10*). After the purge is complete, wake up the system and switch on both the electron beam (EB) and ion beam (IB). Confirm that the “touch alarm is enabled. Apply a 180° scan rotation to both the electron beam and ion beam, select the grid holder, and move the stage to the predefined “Mapping” position.
2. Acquire an electron-beam overview of the grid.
3. At low magnification, center the grid and focus the electron beam (EB). Then, increase the magnification until a single grid square is visible, refocus the EB, and link the Z height by selecting “Link Z to FWD.”
4. Create a new project in the acquisition software and acquire an EB overview image of the grid by creating and naming a new Layer and a new Tileset.
5. Define the Tileset parameters and begin acquisition of the overview image (Table 1).
6. Go to the predefined “Deposition” position and deposit a protective layer of organometallic platinum using the gas injection system (GIS). In the cryo GIS deposition tab, select the grid and set the deposition time to 120–130 s to coat the region prior to milling (Figure 1B) (*see Note 11*).
7. Return to the predefined “Mapping” position and define regions of interest (ROIs) and preliminary lamella sites. With the touch alarm enabled, move the stage to orient the grid orthogonal to the ion beam (IB). For a standard TFS Aquilos 2 shuttle with a 25° pretilt, rotate the stage to 108.1° (180° relative to the Mapping position) and tilt to 7°, positioning the ion beam perpendicular (90°) to the sample surface. Save this shuttle position for future navigation.
8. Wake up the system (*see Note 12*). Set the ion beam (IB) current to 10 pA, navigate to a preliminary lamella site, and adjust image contrast. Acquire an image while focusing the IB during acquisition to minimize charging effects.
9. Confirm that the sample surface is flat and free of contamination. If grid bars are not visible, mill small windows at 15 nA in locations where grid bars are expected (Figure 2A). If orthogonal grid bars are identified, mark lines extending across the grid squares that contain candidate lamella sites. For 200-mesh grids with aligned bars, define the recommended precut rectangles (top: 22 µm × 37 µm; bottom: 20 µm × 17 µm; spacing: 25 µm) and mill these regions at 15 nA (Figure 2B). After completing the precuts, switch the ion beam current back to 10 pA, move to the next preliminary lamella site, and repeat the imaging, surface inspection, grid bar identification, and precutting steps as needed.
10. After completing the precuts, return to the “Mapping” position and acquire a new Tileset using the same parameters as previously defined. Delete the old preliminary lamella sites and redefine lamella sites based on the precut positions. Open AutoTEM, select the project with the same Project Name used in MAPS, and choose the appropriate milling template (Table 2). Apply the template to all defined lamella sites and run “Preparation” in Guided mode (Figure 2C). Follow the prompts to perform the eucentric tilt adjustment and set the milling angle to 20°.
11. Use the positioning guides to mark the future notch placement. Perform bulk material removal by first defining a rectangular milling pattern at 10 pA, then milling at 7 nA until the bottom bulk material is cleared. Next, tilt the stage to the final milling angle (20°). Define a second rectangular milling pattern at 10 pA, position it a few microns from the slab edge (Figure 2D), and mill at 3 nA until the excess material is removed. This pre-cut cleanup procedure was then repeated at stage tilt angles of 30° and 41° to ensure complete removal of excess material and achieve fully cleaned pre-cuts.
12. For notch milling, align the notch pattern approximately with the initial AutoTEM lamella position; precise alignment is not required at this stage. Define the notch geometry (∼3.5 µm × ∼8.1 µm × ∼3 µm) using the specified sub-pattern sizes, overlaps, and spacing (Figure 2E). Mill the notch at 0.3 nA for approximately 2 min 15 s, adjusting the milling time as needed for different milling angles (Figure 2F).
13. Rerun Image Acquisition and Lamella Placement, then position the lamella approximately 1 µm from the notch with the top and bottom milling boxes overlapping the notch region.
14. Define the final lamella size for milling. Lamella dimensions of ∼12 µm × ∼15–20 µm × ∼150–200 nm are suitable for most samples (Figure 2G-I).
15. Run the workflow in stepwise mode according to the selected AutoTEM template (Table 2) (*see Note 13*).

**Figure 2.**
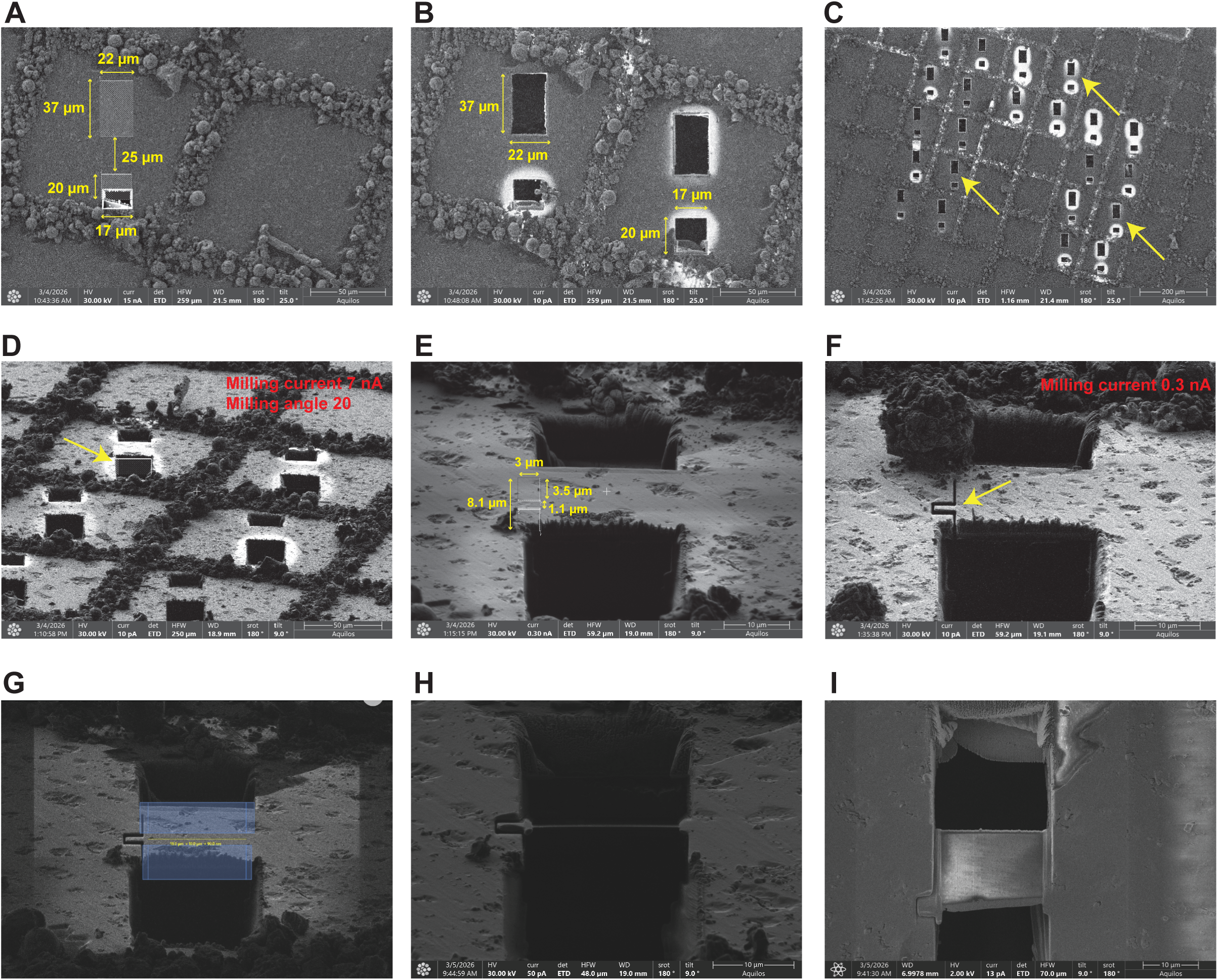
Representative images showing different stages of cryo-FIB milling. **(A)** Small milling windows are generated to identify the positions of underlying grid bars. The yellow arrows show the dimensions and spacing used to determine the milling area. **(B)** Precut milling is performed around the candidate lamella region at high current. The yellow arrows indicate the dimensions and spacing of the precut regions. **(C)** A representative image of the ion beam view of multiple lamella sites after importing them into AutoTEM. The yellow arrows indicate representative sites selected for automated milling. **(D)** The first pre-cut cleanup step is performed at a 20° milling angle with an ion beam current of 7 nA. The yellow arrow indicates the milling region used to remove excess material adjacent to the slab edge. The pre-cut cleanup procedure is repeated at stage tilt angles of 30° and 41° to facilitate the complete removal of residual material. **(E)** The notch-milling pattern region is positioned adjacent to the lamella site. The measurements demonstrate the notch dimensions and spacing parameters. **(F)** Notch pattern after milling at 0.3 nA. The yellow arrow shows the completed notch adjacent to the lamella site. **(G)** Lamella placement in AutoTEM, with upper and lower milling boxes positioned to overlap the notch region. The regions covered with blue boxes will be milled to generate the final lamella with the dimensions of 18 μm x 10 μm x 90 nm. **(H)** A representative ion beam image showing the lamella and the notch after automated milling. **(I)** A representative electron beam image of the final lamella from a different angle.

**Table 1.** Parameters for the EB grid overview acquisition in MAPS.

|  |  |
| --- | --- |
| **Acquisition Type** | Electron |
| **Tiles X, Y** | 5, 7 |
| **Tile HFW** | 600 $\mu\text{m}$ |
| **Total HFW** | 2.76 mm |
| **Resolution** | 3071 x 2048 |
| **Pixel Size** | 195.3125 nm |
| **Dwell** | 1 $\mu\text{s}$ |
| **Frames** | Check, 1 |
| **Reduced Area** | Uncheck |

**Table 2.** Cryo-FIB milling parameters in AutoTEM.

|  | **Lamella thickness**<br>**($\mu\text{m}$)\*** | **Depth Correction**<br>**(%)** | **Front Width Overlap**<br>**($\mu\text{m}$)** | **Rear Width Overlap**<br>**($\mu\text{m}$)** | **Milling Current**<br>**(nA)** | **Pattern Overlap**<br>**(%)** | **Pattern Type** | **DCM Interval**<br>**(s)** |
| --- | --- | --- | --- | --- | --- | --- | --- | --- |
| **Rough Milling** | 1 | 25 | 1.5 | 1 | 1 | N/A | Rectangular | 120 |
| **Medium Milling** | 0.8 | 100 | 0.65 | 0.5 | 1 | 400 | CCS | 90 |
| **Fine Milling** | 0.6 | 100 | 0.35 | 0.1 | 0.5 | 200 | CCS | 60 |
| **Finer Milling** | 0.4 | 100 | 0.05 | 0.05 | 0.3 | 200 | CCS | 30 |
| **Polishing 1** | 0.15 | 75 | N/A | N/A | 0.1 | 300 | CCS | 300 |
| **Polishing 2** | 0 | 30 | N/A | N/A | 0.05 | 400 | CCS | 150 |

\*Lamella thickness is defined as “Pattern Offset”.

### 3.5 Image acquisition

Cryo-ET data acquisition is initiated by defining the target resolution and selecting imaging parameters, while considering both the desired level of detail and the specimen’s structural characteristics. Image acquisition parameters are often refined iteratively with the guidance of tilt-series alignment accuracy, preprocessing parameters, and subtomogram averaging (STA) quality. Precautions are taken to minimize radiation damage during target identification, centering, and eucentric height adjustment, as well as to reduce specimen drift during exposure. Image acquisition in movie mode allows for the subsequent motion correction process to compensate for residual drift. To maintain accurate focus throughout the tilt series, off-target focus areas are used. Additionally, ensuring the parallel illumination of a highly coherent 300 keV electron beam is critical for preserving high-resolution information.

1. Load vitrified grids containing cryo-FIB milled lamellae into the Krios cryo-TEM and adjust microscope parameters before data acquisition (*see Note 14*).
2. In SerialEM (v4.1.0) (Mastronarde, 2005), create a whole-grid atlas at low magnification (125x) to assess grid quality, ice thickness gradient, and lamella integrity *(see Note 15)*.
3. Acquire square montages (medium magnification) for optimal regions.
4. Identify the lamellae suitable for tomography based on ice thickness, grid contamination, and intact lamella geometry.
5. Add Navigator points on selected lamella regions.
6. Move the stage to each selected Navigator point.
7. Acquire a low-dose preview image for each lamella target and optimize target positioning by centering the region of interest within the lamella.
8. Save this image as the Anchor map.
9. Repeat anchor map generation for all selected targets using: SerialEM → “Anchor Map” button (*see Note 16*).
10. Set the record area to match the acquisition magnification and pixel size for each data-collection condition separately (*see Note 17*).
11. Define a range of defocus values (−2 µm to −5 µm) across targets in 0.25 µm increments (*see Note 18)*.
12. Configure tilt series in SerialEM using the Tilt Series Setup/Start (TS) module. Employ a dose-symmetric tilt scheme, beginning at 0° relative to the lamella pre-tilt, with a tilt increment of 2° between ±60°.
13. Acquire data in groups of two tilts per side, following a dose-symmetric acquisition pattern.
14. Record images with SerialEM at a nominal magnification of 64,000x, corresponding to a pixel size of 1.965 Å. The dose rate per tilt angle will be set at ∼2 e⁻/Å², adding up to a total accumulated dose of ∼100 e⁻/Å² for each tilt series (*see Note 19)*.
15. Employ standard tomography tracking and autofocus tools in SerialEM during tilt-series acquisition (*see Note 20*).
16. Perform automated acquisition in SerialEM using the Navigator. Mark targets as individual acquisition points, designate for tilt-series collection by enabling the “TS” flag, and sequentially collect using the “Acquire at Items” function.
17. Configure acquisition to use the Falcon 4i in full-frame mode at 4096 × 4096 pixels, with the detector operating in counting mode and saving data in EER format.

### 3.6 Image processing

Cryo-ET data processing largely follows principles established for STA workflows, with additional optimization steps to enhance tomogram quality, signal-to-noise ratio, and downstream interpretability. In practice, achieving high-resolution structures requires coordinated use of multiple specialized software packages along with custom scripts that bridge processing steps and create an integrative platform.

Accordingly, this section describes a data processing strategy for analyzing raw tilt-series using an integrated workflow that combines *IMOD* for stack assembly, visualization, and tilt-series handling (Mastronarde and Held, 2017); *AreTomo3* for motion correction, CTF estimation and correction, tilt-series alignment, tilt-axis refinement, and weighted back-projection reconstruction (Peck et al., 2025a); *IsoNet2* for deep learning–based denoising and missing-wedge compensation (Liu et al., 2025); and *MemBrain v2* for automated membrane segmentation and feature extraction (Lamm et al., 2025) (Figure 3).

**Figure 3.**
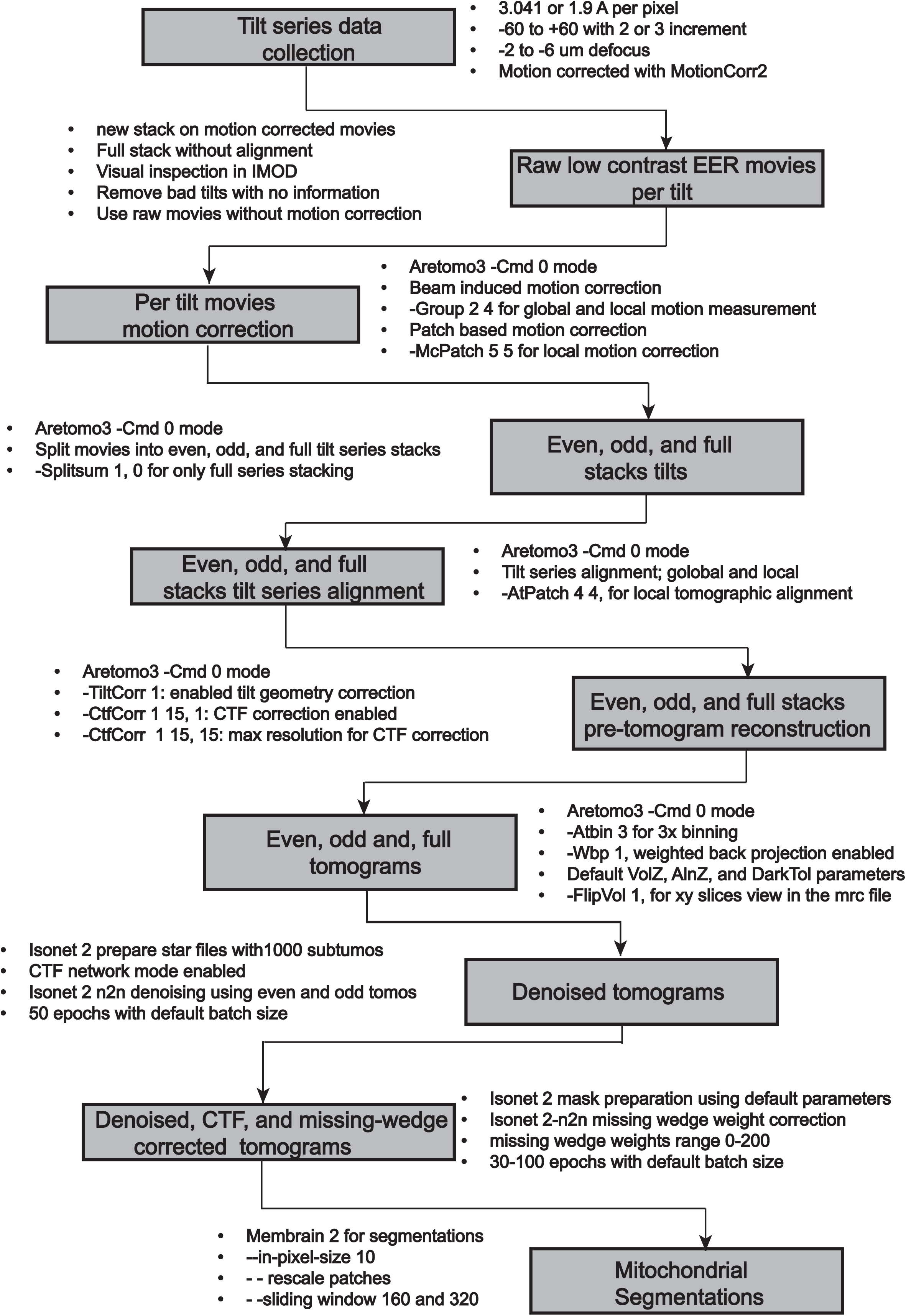
Cryo-ET image processing workflow. A diagram summarizing the data acquisition and image processing steps and parameters for cryo-ET. Motion-corrected tilt images are assembled and inspected in IMOD to identify and eliminate low-quality tilts. The corresponding raw EER movies are processed in *AreTomo3* Cmd 0 mode for per-tilt motion correction, generation of even, odd, and full tilt-series stacks, tilt-series alignment, tilt-geometry correction, CTF correction, and weighted back-projection reconstruction. Even and odd tomograms are imported into *IsoNet2* for STAR generation, *noise2noise* denoising, mask generation, and missing-wedge correction. The final denoised, CTF-corrected, and missing-wedge-corrected tomograms are loaded into *MemBrain v2* for segmentation.

We applied these tools sequentially within a unified workflow, with parameters carefully optimized at each stage to preserve data integrity while improving signal-to-noise ratio and segmentation accuracy. This modular strategy enables systematic refinement of reconstruction, denoising, and segmentation steps, ensuring consistent processing across datasets while retaining flexibility for dataset-specific optimization.

#### Steps for Tilt-Series (Stack) Assembly

1. Convert motion-corrected tilt images (*see Note 21)* and their metadata into an ordered tilt-series stack suitable for downstream alignment and tomographic reconstruction. Because the motion-corrected projections are produced as individual files, a custom script termed `MC stack + tilt map` is used to reconstruct their original acquisition order, record tilt metadata, and assemble a reconstruction-ready stack using the steps below (Figure 3 and Code 1).
2. Run the `MC stack + tilt map` on the acquisition output directory containing the motion-corrected tilt images `(*.mc.mrc)` and metadata files `(*.mdoc)`from the data collection session (see *Note 22)*.
3. Check the output stack against `sorted_filelist.txt` to confirm that the number of sections matches the number of listed tilt images (see *Note 23)*.
4. Remove tilt images unsuitable for downstream reconstruction while preserving the identities of the retained tilts. Because tilt exclusion must be propagated back to the original raw data, a custom cleaning script is used to map excluded views from the motion-corrected tilt-series stack to the corresponding raw tilt files (Figure 3 and Code 2).

#### Steps for Stack Cleaning

5. Open the full motion-corrected tilt-series stack in IMOD/3dmod and visually inspect the stack to identify unusable tilts (e.g. those affected by contamination, severe drift, tracking loss, or absence of interpretable signal).
6. Provide the indices of the tilts to be excluded as input to the custom cleaning script (*see Note 24*). The script uses this information to identify the corresponding raw tilt files, remove the excluded tilts from further processing, and generate a cleaned tilt-angle list together with a `KEEP_INDEX` file that preserves the original acquisition indices of the retained tilts (Code 2).
7. Define the cleaned reconstruction input for the subsequent processing stage. The `KEEP_INDEX` file preserves the identities and original ordering of the retained tilts, thereby enabling reproducible downstream reconstruction.

#### Steps for Motion Correction, Alignment, and Reconstruction

8. Use the retained raw tilt files together with the cleaned tilt-angle list from the stack-cleaning step as input for tomographic reconstruction. *AreTomo3* is an integrated software package for automated marker-free, motion-corrected cryo-electron tomographic alignment and reconstruction (*see Note 25)*.
9. In a Linux terminal on a GPU node, run *AreTomo3* `Cmd 0` to systematically perform motion correction, tilt-series alignment, CTF correction, and 3D tomographic reconstruction (Figure 4) (*see Note 26*).

~~~
AreTomo3 -InPrefix <input_prefix> -InSuffix <.mdoc> -OutDir
<output_directory> -Gain <gain_reference.mrc> -FmIntFile
<frame_interval_file> -PixSize <pixel_size> -Kv <300> -Cs <2.70> -Group <2
4> -McPatch <5 5> -AtPatch <4 4> -TiltCorr <1> -CtfCorr <1 15> -Wbp <1> -
AtBin <3> -SplitSum <1> -FlipVol <1> -Gpu <gpuid>
~~~

**Figure 4.**
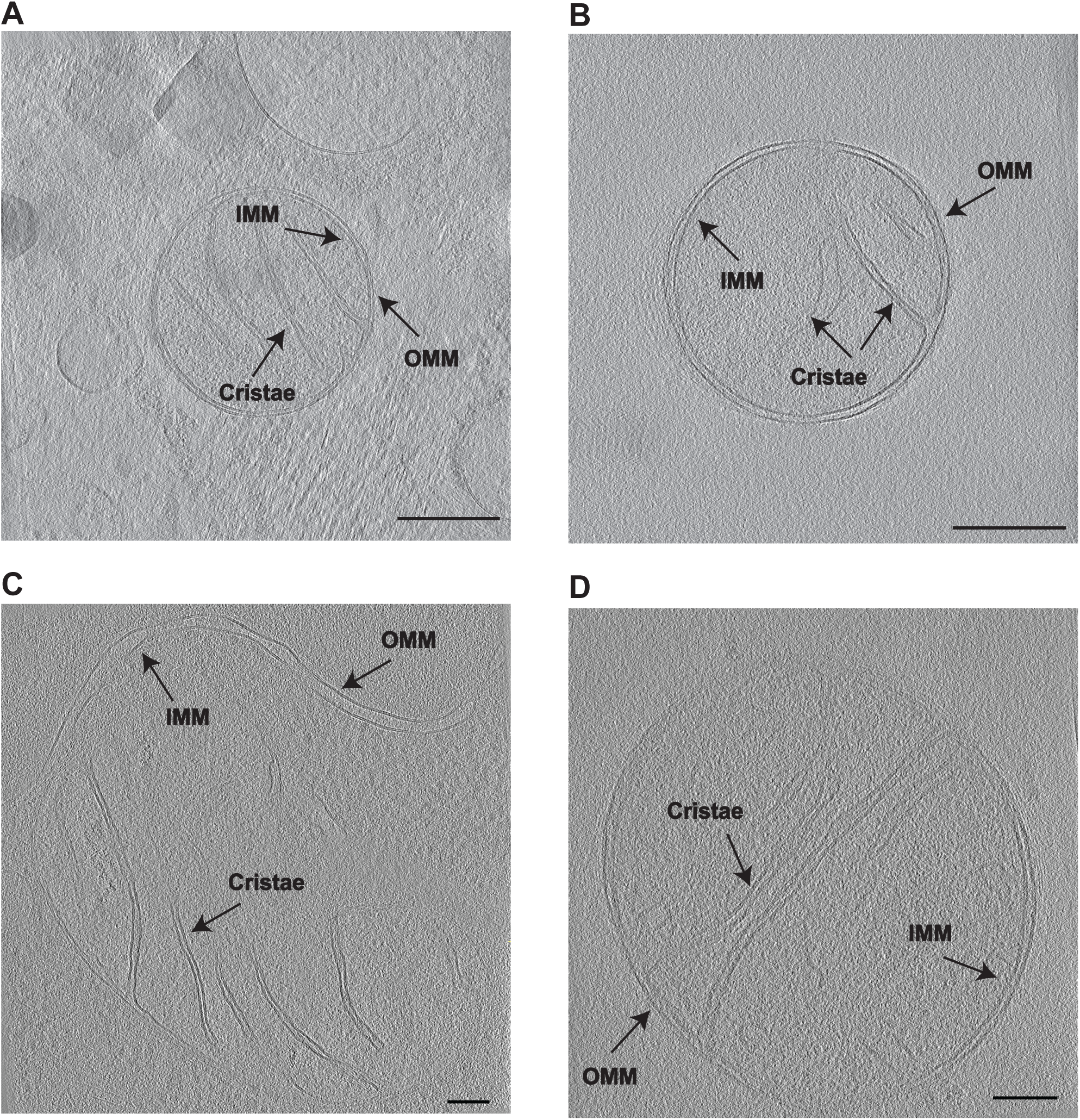
Representative reconstructed tomograms of wild-type mitochondria isolated from human HAP1 cells. (**A-D)** Slices from AreTomo3-reconstructed cryo-electron tomograms illustrate mitochondrial ultrastructure, including the outer mitochondrial membrane (OMM), inner mitochondrial membrane (IMM), and cristae, as indicated. The panels display representative examples of various mitochondrial morphologies resolved after motion correction, tilt-series alignment, CTF correction, and weighted back-projection reconstruction. Scale bars, 25 nm (A, B) and 10 nm (C, D).

#### Steps for denoising and Missing-Wedge Refinement

10. Perform denoising and missing-wedge refinement after tomographic reconstruction to improve interpretability and recover structural information lost during limited-angle data collection.
11. Generate paired even and odd tomograms from *AreTomo3*, which provide the independent inputs required for *IsoNet2 Noise2Noise* denoising.
12. Prepare a *tomograms.star* file to define the tomogram paths and acquisition parameters needed for downstream processing.
13. Use the denoised tomograms to generate masks that exclude empty or uninformative regions, improving the efficiency and accuracy of missing-wedge correction.
14. Perform *IsoNet2* refinement using the denoised, masked tomograms to reduce missing-wedge artifacts and produce corrected volumes for visualization and analysis (*see Note 27*).
15. Confirm that even and odd tomograms are produced from *Aretomo3* reconstruction.
16. Load *isonet.py* in a Linux terminal to execute *prepare_star module* according to the described usage (*isonet.py prepare_star --help*). The parameters provided below are an example and should be optimized for each dataset (Figure 5).

~~~
isonet.py prepare_star --even_dir <even_tomogram_directory> --odd_dir
<odd_tomogram_directory> --pixel_size <pixel_size_a_per_pix> --defocus
<defocus_in_a> --tilt_min <tilt_min_angle> --tilt_max <tilt_max_angle> --
number_subtomos <1000> --output_star <tomograms.star>
~~~

17. Load *isonet.py* to execute *Noise2Noise (n2n)* `denoise` module (Figure 3) (*see Note 28*).

**Figure 5.**
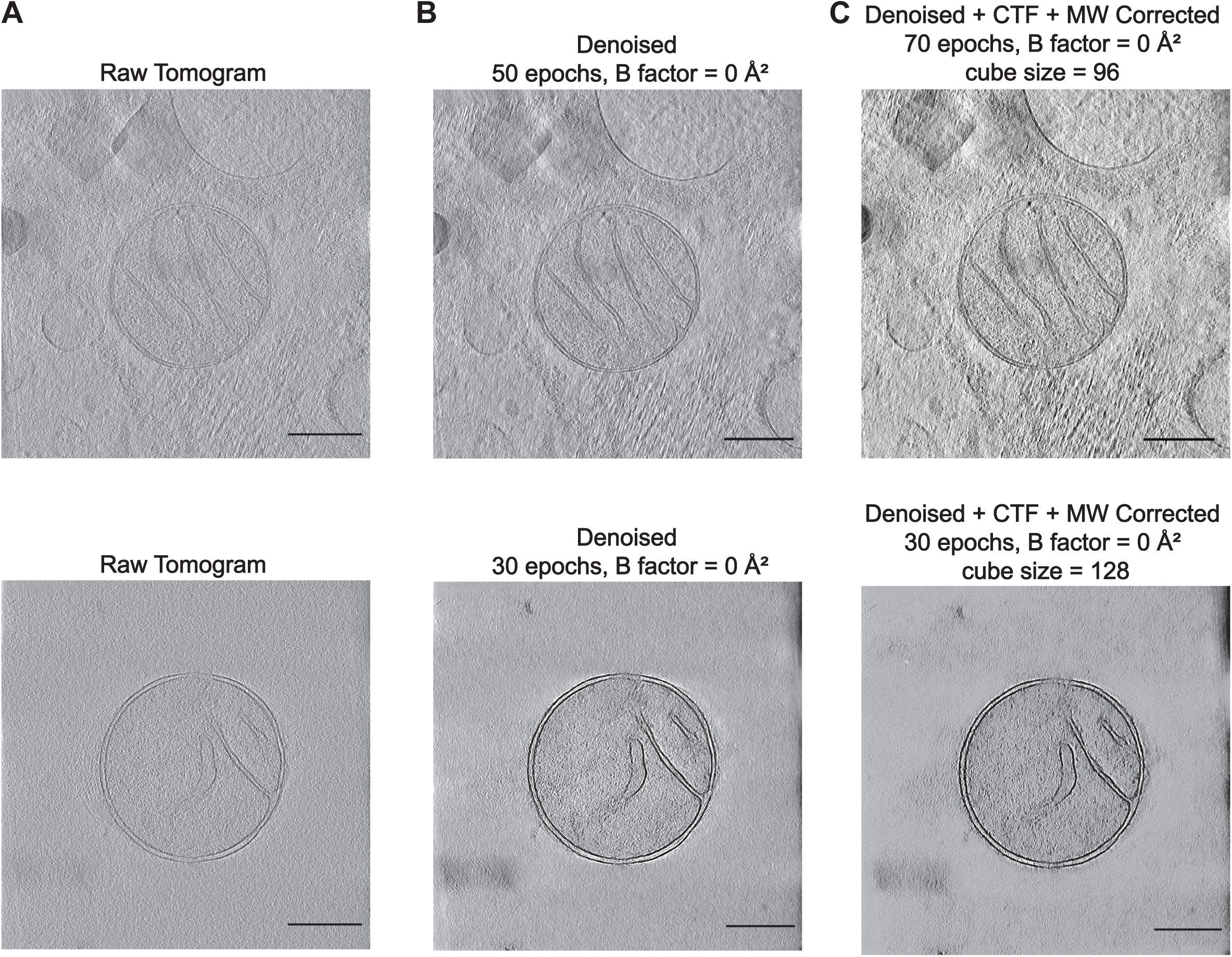
Representative XY-view slices of original, *IsoNet2*-denoised, and missing-wedge-corrected tomograms obtained using different refinement and mask-generation parameters. **(A)** Raw reconstructed tomogram slices (top and bottom). **(B)** Denoised tomograms generated using the raw tomograms in panel 5A as input. The *IsoNet2 Noise2Noise* module was used to denoise even and odd reconstructions as independent noisy observations, with the parameter CTF_mode = network enabled in all cases. Only the number of training epochs and the B factor were varied between examples, while all other denoising parameters were kept at their default settings. The denoising settings were 50 epochs, B factor = 0 Å² (top), 30 epochs, B factor = 0 Å² (bottom). **(C)** Missing-wedge refinement was performed in *IsoNet2* using the denoised tomograms as input, with masks generated from the denoised tomograms before refinement. The missing-wedge correction and mask-generation parameters were varied across examples as follows: 70 epochs, B factor = 100 Å², density_percentage = 50, std_percentage = 80, cube size = 96 (top), 30 epochs, B factor = 0 Å², density_percentage = 50, std_percentage = 50, cube size = 128 (bottom). Scale bars, 25 nm.

~~~
isonet.py denoise <STAR_FILE> --output_dir <output_dir> --arch <unet-
medium> --cube_size <cube_size> --epochs <epochs> --loss_func <L2> --
save_interval <save_interval> --learning_rate <learning_rate> --
learning_rate_min <learning_rate_min> --mixed_precision <True|False> --
CTF_mode <network|phaseflip|none> --isCTFFlipped <True|False> --
do_phaseflip_input <True|False> --bfactor <bfactor> --clip_first_peak_mode
<clip_first_peak_mode> --snrfalloff <snrfalloff> --deconvstrength
<deconvstrength> --highpassnyquist <highpassnyquist> --with_preview
<True|False> --gpuID <gpuID>
~~~

18. Load *isonet.py* to execute `make_mask` module to generate mask using the input STAR file containing the denoised tomogram paths for excluding empty, non-informative regions before missing-wedge correction (Figure 3) (*see Note 29*).
19. Open the denoised tomogram and the corresponding mask in *IMOD* and visually inspect the mask to confirm appropriate coverage of informative regions and exclusion of empty space.

~~~
isonet.py make_mask <STAR_FILE> --output_dir <output_dir> --
density_percentage <density_percentage> --std_percentage <std_percentage>
--patch_size <patch_size> --z_crop <z_crop> --use_gpu <True|False>
~~~

20. Run *isonet.py* `refine` to perform missing-wedge correction (Figure 6) on the denoised tomograms using the prepared `tomograms.star` file as input. In this step, the denoised tomograms (*rlnDenoisedTomoName*) are used as the refinement source. If mask generation was performed previously, the corresponding masks are referenced through the STAR file and applied during refinement (Figure 3). The parameters provided below reflect the configuration used and may require optimization for each dataset. Visualize the missing wedge corrected tomogram in *IMOD* (Figure 5).

**Figure 6.**
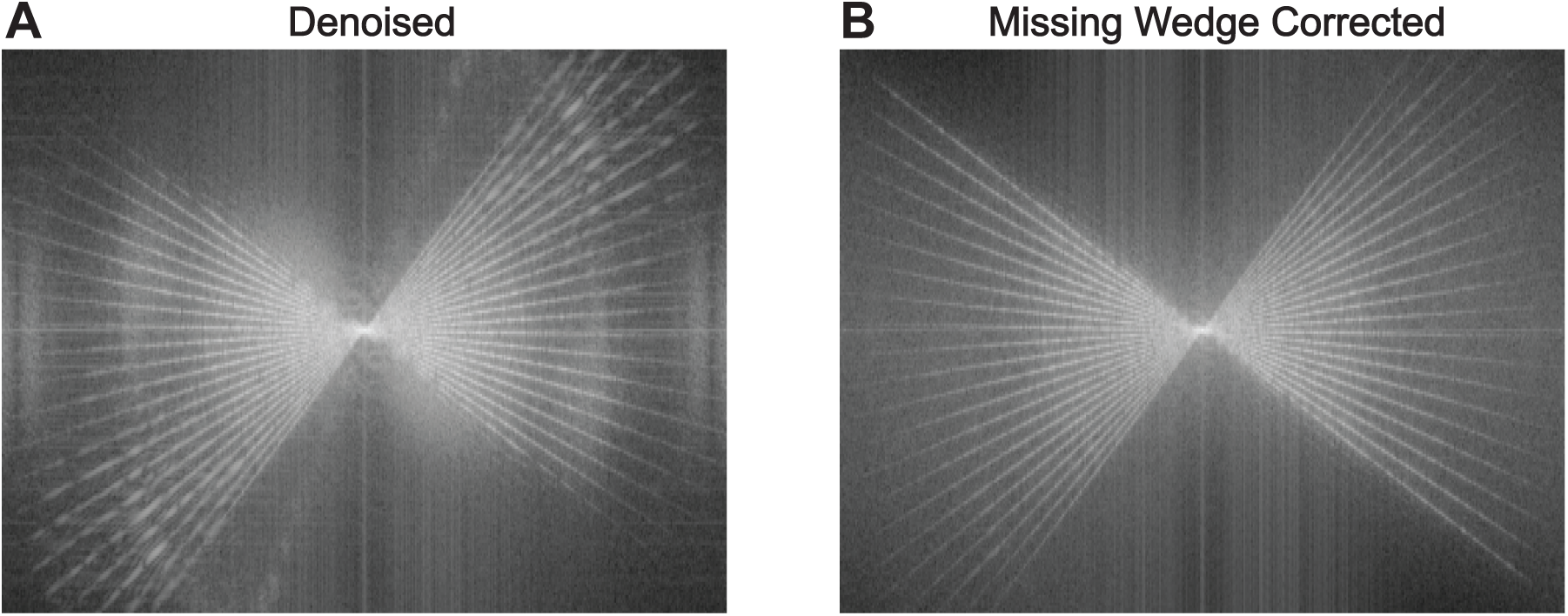
Denoising and missing wedge correction of tomograms. **(A)** Power spectrum of a representative denoised tomogram. **(B)** Power spectrum after missing-wedge correction of the denoised tomogram shown in panel 6A.

~~~
isonet.py refine <STAR_FILE> --output_dir <isonet_maps> --method <isonet2-
n2n> --input_column <rlnDenoisedTomoName> --arch <unet-medium> --cube_size
<128> --epochs <100> --batch_size <auto> --loss_func <L2> --learning_rate
<0.0003> --learning_rate_min <0.0003> --save_interval <10> --mw_weight <20>
--apply_mw_x1 <True> --mixed_precision <True> --CTF_mode <network> --
clip_first_peak_mode <1> --bfactor <0> --isCTFFlipped <False> --
do_phaseflip_input <True> --noise_level <0> --noise_mode <nofilter> --
random_rot_weight <0.2> --with_preview <True> --snrfalloff <0> --
deconvstrength <1> --highpassnyquist <0.02> --gpuID <gpuID>
~~~

#### Steps for tomogram segmentation

21. In a Linux terminal, load *MemBrain v2* software to perform tomogram Segmentation (*see Note 30*).
22. Execute `tomo_preprocessing <command> --help` to list the available preprocessing modules and their specific flags.
23. Execute `match_pixel_size` to resample tomograms to the pixel size required for downstream MemBrain processing. The parameters provided below are an example and should be optimized for each dataset.

~~~
tomo_preprocessing match_pixel_size --input-tomogram <PATH> --output-path
<path> --pixel-size-out 10.0 --pixel-size-in <size>
~~~

24. Run `membrain segment` on the denoised and missing-wedge-corrected tomogram to perform membrane segmentation, using the trained *MemBrain-v2* model checkpoint specified by `--ckpt-`path.
25. Specify the tomogram to be segmented with --tomogram-path and the trained *MemBrain-v2* model checkpoint with `--ckpt-path`.
26. Specify the output directory with `--out-folder`.
27. Use `--rescale-patches` to rescale the tomogram to 10 Å when required.
28. Use `--sliding-window-size` to define the prediction window size and `--in-pixel-size` to manually specify the input pixel size if the tomogram header is incorrect. The parameters provided below should be optimized for each dataset (Figure 7).

**Figure 7.**
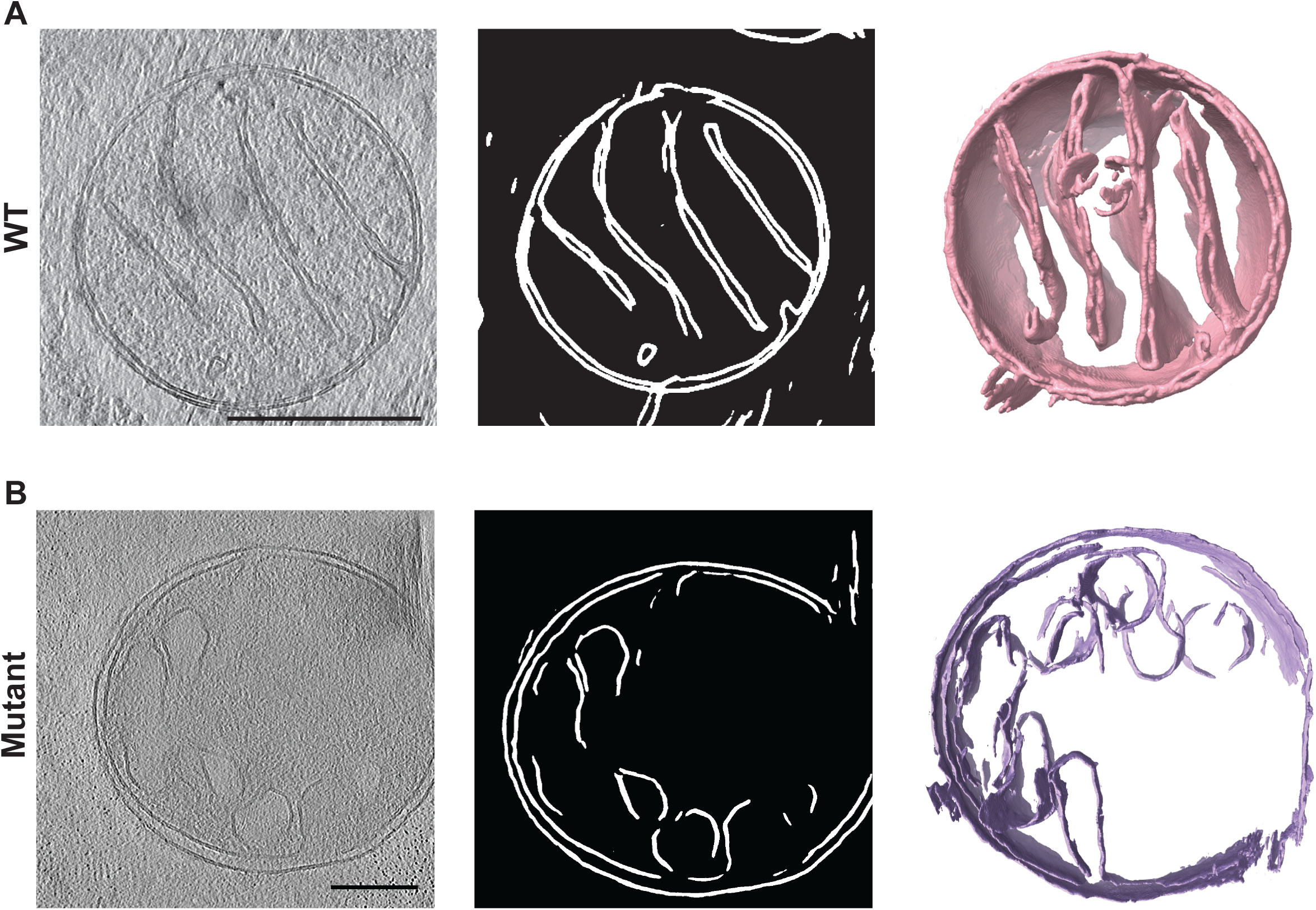
Automated segmentation of denoised and missing-wedge-corrected tomograms. **(A, B)** Representative images of the input tomogram slice (left), the corresponding segmented view of a slice (center), and the rendered segmented volume (right) are shown WT (A) and mutant mitochondria (B). Automated membrane segmentation was performed using MemBrain v2 in both cases, with the parameters sliding-window-size = 320 in (A) and sliding-window-size = 160 in (B). Scale bars, 25 nm.

~~~
membrain segment --tomogram-path <path> --ckpt-path <model_path> --out-
folder <directory> --rescale-patches --sliding-window-size <int> --in-
pixel-size <float<
~~~

29. Open the segmented volume in *ChimeraX* to visualize the structural features (Figure 7).

Defining molecular resolution features of key cellular machines, including mitochondrial OXPHOS complexes and ribosomes, typically require collecting several tilt series and subsequent subtomogram averaging (STA) and alignment data processing steps (Anton et al., 2025; Zheng et al., 2024). Additionally, data acquisition at higher magnification with a smaller calibrated pixel size and further optimization of imaging conditions is recommended for capturing high-frequency information and determining the 3D reconstructions of proteins and protein complexes at higher resolution (*see Note 31)*. High-quality subtomograms can be used for model building and refinement by leveraging recent advancements in structure prediction and model building tools (Adams et al., 2010; Baek et al., 2021; Emsley et al., 2010; Fadini et al., 2025; Jamali et al., 2024; Jumper et al., 2021).

## 4. Results

We established an integrated workflow capable of generating high-quality 3D reconstructions of human mitochondria while maintaining their native structural state. By optimizing the most labor-intensive stages of data acquisition and image processing, we significantly streamlined the experimental workflow without compromising the structural integrity of the mitochondria in the resulting 3D volumes. These results demonstrate that the optimized pipeline provides a high-performance framework for investigating mitochondrial dynamics across both healthy and pathological states.

Mitochondria were fluorescently labeled and isolated from human HAP1 cells. To preserve the organelle’s structural integrity, samples were vitrified using the waffle method combined with high-pressure freezing (HPF). We subsequently performed cryo-fluorescence screening to rapidly identify regions containing MitoTracker Green-labeled mitochondria (Figure 1). Targeted cryo-FIB milling was then employed to thin these selected regions into lamellae of optimal thickness and geometry (Figure 2). This robust pipeline successfully produced electron-transparent lamellae suitable for cryo-ET imaging.

We acquired several tilt series from different samples using a TFS Krios cryo-transmission electron microscope (cryo-TEM) operated at 300 KeV and equipped with a Falcon 4i direct electron detector and Selectris energy filter. We processed these tilt series through an integrated computational pipeline (Figure 3), beginning with stack assembly in *IMOD*. We then employed *AreTomo3* for motion correction, CTF estimation and correction, and weighted back-projection reconstruction steps, yielding reconstructed 3D tomograms (Figure 4). To optimize these volumes for downstream analysis, we applied *IsoNet2* for denoising and missing-wedge refinement (Figures 5 and 6). This significantly enhanced the contrast and signal-to-noise ratio, enabling precise membrane segmentation using *MemBrain v2* (Figure 7).

We applied this workflow to the 3D visualization of mitochondrial ultrastructure and the characterization of membrane-associated features. Comparative analysis of reconstructed and segmented tomograms from wild-type (WT) and mutant samples demonstrated that mutations within the OXPHOS machinery severely disrupt cristae architecture (Figure 7). These findings underscore the utility of this integrated platform for investigating the mechanisms of mitochondrial membrane dynamics and understanding how pathologies associated with mitochondrial dysfunction influence organelle ultrastructure.

## 5. Notes

1. The optimal number of homogenization strokes required for effective cell disruption varies depending on cell type and cell density. This parameter should be empirically determined for each experiment by periodically examining the homogenate under a standard cell culture microscope to assess the extent of cell lysis. Additionally, different motors with speed control can be used for homogenizing the sample.
2. Working with liquid nitrogen is a safety hazard and must be performed using appropriate precautions. Always wear proper personal protective equipment (PPE), including cryogenic gloves, safety goggles or a face shield, and ensure the use of oxygen monitoring systems in enclosed or poorly ventilated areas.
3. Proper grid orientation during clipping is critical to ensure that lamellae can be accurately positioned for subsequent cryo-FIB milling and cryo-electron tomography data acquisition.
4. The additional carbon layer enhances grid rigidity, improving structural stability during high-pressure freezing and subsequent handling steps.
5. In addition to the TEM grid type used in this study, other standard grid types with different materials and mesh sizes can be utilized for sample preparation. While the mesh size is an important factor for waffle formation and handling, the grid material is generally less critical.
6. If excess sample is present during planchette assembly, gently blot to remove surplus liquid while ensuring the formation of a uniform waffle.
7. In most cases, the waffle assembly should detach cleanly from the planchettes. If the planchettes remain adhered, gently separate them by applying slight pressure with tweezers; a second pair of tweezers may be used to stabilize the assembly. If only one planchette separates, carefully grasp the grid at the outer rim and lift it away from the remaining planchette.
8. The clipping process must be performed under cryogenic conditions to preserve vitrification. First, transfer the vitrified grid to a liquid nitrogen-cooled workstation. Place the AutoGrid base in the clipping station and carefully position the grid inside it using pre-cooled, fine-tipped tweezers. Under a stereomicroscope, rotate the grid so that the grid bars are aligned with the AutoGrid notch, which defines the orientation for downstream milling and data collection.
9. After proper alignment, gently press a C-clip ring into place to secure the grid within the AutoGrid. Minor movement during clipping is acceptable, provided overall alignment is maintained. The clipped grid is then ready for transfer into cryo-FIB/SEM for lamella preparation and subsequent tomography.
10. Ensure that no sample is present during the GIS purge to prevent contamination of the injection system and maintain stable deposition performance.
11. Conductive platinum sputter coating is omitted in this workflow, as the organometallic platinum layer deposited using the GIS provides sufficient surface protection and minimizes charging during cryo-FIB milling.
12. The system may automatically enter standby after sputtering; ensure it is fully reactivated and stable before proceeding with imaging or milling.
13. Perform optional final polishing or over-tilt milling, depending on lamella thickness and quality. If the lamella or GIS platinum layer appears damaged, skip the automated post-milling step.
14. Prior to data acquisition, ensure the microscope is properly aligned and stable. This includes confirming stable emission from the E-CFEG source, aligning the beam for parallel illumination, inserting the Selectris X energy filter with a 10 eV slit width, and verifying that the Falcon 4i detector is active and properly configured. Additional alignments, such as coma-free alignment and astigmatism correction, should be performed to ensure optimal imaging conditions.
15. In addition to the SerialEM (v4.1.0) software (Mastronarde, 2005), Thermo Fisher Scientific provides Tomography 5 software to support flexible cryo-ET workflows. In this study, we utilized SerialEM as it is a well-established and widely adopted cryo-ET data acquisition platform that provides scripting capabilities needed for our data collection strategy.
16. Anchor maps are used for target tracking during tilt-series acquisition and for realignment between tilts, ensuring consistent positioning of the region of interest throughout the tilt series.
17. To avoid mixing acquisition conditions, save each imaging configuration as a separate SerialEM imaging state or settings file (e.g., 42k and 64k). For the 42k condition, data are acquired at a nominal magnification of 42,000× (3.041 Å/pixel), while for the 64k condition, data are acquired at 64,000× (1.965 Å/pixel). In both cases, images are recorded using a Falcon 4i direct electron detector in counting mode, with full-frame acquisition (4096 × 4096 pixels) and saved in Electron Event Representation (`.eer`) format.
18. Defocus values are selected to balance phase contrast and high-frequency signal. Increasing defocus enhances image contrast, which facilitates tilt-series alignment and target tracking, but reduces high-resolution information. Therefore, defocus values are systematically varied across targets within a range of −2 µm to −5 µm in 0.25 µm increments to optimize both contrast for alignment and resolution for downstream 3D reconstruction and subtomogram averaging (STA) steps.
19. The nominal magnification and calibrated pixel size should be determined by the sample type and the target Nyquist resolution. Therefore, the optimal pixel size is determined empirically to balance resolution requirements with downstream data processing strategies. The total accumulated dose per tilt series, which is typically maintained around 100 to 120 e⁻/Å², is user-defined and may vary depending on the acquisition settings, tilt schemes, and sample-specific imaging requirements.
20. Target tracking is performed using low-dose trial images, while autofocus is carried out in a designated off-target focus area to minimize radiation damage to the region of interest. Alignment between tilts is maintained using previously acquired anchor maps. Trial and focus areas are positioned along the tilt axis, and beam conditions are kept consistent throughout the tilt series to ensure stable tracking and imaging.
21. The motion-corrected tilt images generated during data collection are first used for stack assembly, visual inspection, and removal of unsuitable tilts. This screening step is performed on the motion-corrected projections because they allow rapid evaluation of tilt quality while preserving traceability to the original raw data. After unsuitable tilts are removed, the corresponding retained raw (`.eer`) files are used as input for *AreTomo3* `Cmd 0`, which performs motion correction, tilt-series alignment, CTF correction, tomographic reconstruction, and generates even and odd split-sum reconstructions for downstream analysis (Peck et al., 2025a).
22. The script parses each `.mdoc` file to extract the acquisition index and nominal tilt angle for each tilt image, then sorts the images by acquisition index to preserve the original tilt geometry. It generates an ordered image list (`sorted_filelist.txt`) containing one tilt image per line and a tilt metadata table (`tilt_map.tsv`) that records the acquisition index, nominal tilt angle, and corresponding image file. The ordered image list is then used to assemble the tilt images into a single MRC tilt-series stack using IMOD `newstack` (Mastronarde and Held, 2017).
23. The validated stack defines the ordered, motion-corrected tilt series used for inspection and curation prior to reconstruction. This stack serves as the input for tilt cleaning, where unusable tilts are identified and excluded. The associated `sorted_filelist.txt` and `tilt_map.tsv` files provide traceability, allowing each retained or excluded tilt to be linked back to its original acquisition.
24. The cleaning script retrieves the corresponding raw tilt files `(.eer)` from the acquisition directory and excludes specified tilts from further processing. Only retained tilts are propagated for reconstruction. The script generates a cleaned tilt-angle list and a `KEEP_INDEX` file containing the original acquisition indices of retained tilts, and verifies consistency between the retained raw files, tilt angles, and index mapping.
25. In addition to *AreTomo3*, marker-free tilt-series alignment and tomographic reconstruction can also be performed using the *eTomo* (IMOD) (Mastronarde and Held, 2017) and nextPYP (Liu et al., 2023) software packages. Alternatively, EMAN2 (Galaz-Montoya et al., 2015) and Dynamo (Coray et al., 2024) software, which are also widely used for cryo-ET workflows, can perform fiducial-based alignment of tilt series.
26. This step generates full, even, and odd tomograms for downstream denoising and missing-wedge correction. For optimal performance, we recommend working with 3x to 5x downsampled data (corresponding to a pixel size of ∼10 Å).
27. Due to variability in raw tilt-series quality and imaging conditions in each data acquisition, parameters used for `prepare_star`, `denoise`, `make_mask`, and `refine` should be optimized for each dataset.
28. Denoising is performed using *IsoNet2* (Liu et al., 2025) to reduce noise while preserving structural features. The *Noise2Noise* (n2n) framework leverages independent noisy observations (even and odd reconstructions) to train the denoising model. The `CTF_mode=network` option enables the model to account for CTF-induced contrast modulation during training rather than relying on explicit pre-correction.
29. This step improves training efficiency by restricting sampling to informative regions. Parameters such as `density_percentage`, `std_percentage`, and `z_crop` should be optimized for each dataset prior to missing-wedge correction.
30. This segmentation workflow enables rapid identification of prominent mitochondrial features, including the outer and inner mitochondrial membranes and cristae. As an alternative to *MemBrain-v2*, tomogram segmentation can also be performed using semi-automated IMOD/3dmod or EMAN2. Practical protocol for efficient annotation of tomograms using IMOD, EMAN2, and machine learning methods have been described previously (Danita et al., 2022; Peck et al., 2025b).
31. For higher-resolution subtomogram averaging, tomographic data can be further processed using *RELION-5*, which provides an integrated image-processing pipeline for cryo-ET and STA (Burt et al., 2024).

## 6. Discussion

Accumulating evidence suggests that investigating the timing, compartmentalization, communication, and localization of mitochondrial processes is critical for advancing our understanding of cellular physiology, degeneration, and regeneration, and may also offer potential intervention strategies. The dynamics of mitochondrial morphology are a critical determinant of mitochondrial and cellular function. However, the molecular mechanisms of mitochondrial membrane dynamics and how mitochondrial proteins assemble into intricate and precise biomolecular machines are not known. Thus, more quantitative explanations of mitochondrial structure are required to understand the mechanisms that link morphological changes to the organelle’s function. The chief obstacle is to develop physical models that will allow us to make direct observations at molecular resolution. By isolating mitochondria under physiological and pathological conditions and pairing this with correlated light microscopy and high-resolution cryo-ET, we may be able to resolve many questions relating to the key mitochondrial processes that are compositionally complex, dynamic, and challenging to reconstitute. Moreover, this approach will allow us to investigate complex biochemical mechanisms such as the self-assembly, dynamics, and self-repair of OXPHOS complexes and the regulation of mitochondrial membrane dynamics.

The major focus of this protocol is to provide a detailed description of optimal sample preparation and data collection parameters to ensure that high-resolution cryo-ET tilt series can be obtained. The workflow also outlines computational tomogram segmentation steps to illuminate the structural features and spatiotemporal dynamics of human mitochondria at the nanometer scale. The major advantage of this pipeline is that these optimized protocols detail the fundamental steps for any cryo-ET project and can be easily adapted to other studies. Furthermore, our multifaceted approach can be combined with downstream structural analysis, such as subtomogram averaging (STA), and has the potential to determine molecular resolution structures of proteins and protein complexes. Perhaps the most important weakness of the cryo-ET workflow is that it has only been implemented in mitochondria. That said, methods for organelle isolation have been described previously for other subcellular compartments, and similar sample preparation steps can be used to structurally characterize these organelles or to generate thin lamellae of cellular volumes via cryo-FIB milling with minor adjustments.

In summary, our workflow offers cutting-edge sample preparation and image acquisition techniques to obtain high-quality images of human mitochondria and describes image processing strategies for performing segmentation and structural analysis. Together, this optimized imaging and image analysis protocol will enhance our ability to visualize cellular compartments and facilitate the structural characterization of key biomolecular machines.

## Acknowledgments

We would like to thank Dr. Yumeng Liu and Dr. Matt Larson at MCCET for assistance with cryo-ET data acquisition, data transfer, and preprocessing.

## Funding

This work was supported in part by the U.S. Department of Energy Office of Basic Energy Sciences grant under award number DE-SC0025606 (H.A.) and National Institutes of Health grants R35 GM150942 (H.A.) and R35 GM139615 (E.P.). Some of this work was performed at the National Center for In-Situ Tomographic Ultramicroscopy (NCITU) and the Simons Electron Microscopy Center located at the New York Structural Biology Center, supported by the NIH Common Fund Transformative High Resolution Cryo-Electron Microscopy program (U24 GM129539 and U24 GM139171) and by grants from the Simons Foundation (SF349247) and NY State Assembly. Additionally, a portion of this research was supported by NIH grant U24 GM139168 and performed at the Midwest Center for Cryo-ET (MCCET) and the Cryo-EM Research Center in the Department of Biochemistry at the University of Wisconsin-Madison.

## Competing Interests Statement

The authors declare no competing interests.

## Notes

### Competing Interest Statement

The authors have declared no competing interest.

