## Supplementary material for "A robust workflow for 3D imaging of human mitochondria using cryo-electron tomography": Code 1

**Code 1. A framework for preparing full stacks from sorted and motion-corrected tilts**

```
#!/usr/bin/env bash
#
# build_mc_stacks_generic_v3.sh
#
# Purpose:
#   Scan a parent directory for tilt-series folders, find motion-corrected
#   tilt images inside each series, sort them by tilt angle, write a sorted
#   file list, and build one full MRC stack per tilt series using IMOD newstack.
#
# Important:
#   This version does NOT use "newstack -fileinlist".
#   Instead, it reads the sorted file list into a Bash array and passes the
#   files to newstack as positional input arguments, which is often more robust.
#
# Path behavior:
#   - no hard-coded paths
#   - no dependence on current working directory
#   - uses all immediate subdirectories under the chosen parent directory
#
# Expected structure:
#   <PARENT_DIR>/<tilt_series_dir>/<MOTION_SUBDIR>/*_*.mc.mrc
#

set -euo pipefail
IFS=$'\n\t'

echo "=== Build Full Stacks from Motion-Corrected Tilt Images ==="
echo

########################################
# Helper functions
########################################

trim() {
   local s="$1"
   s="${s#"${s%%[![:space:]]*}"}"
   s="${s%"${s##*[![:space:]]}"}"
   printf '%s' "$s"
}

abs_path() {
   local target="$1"
```

```
    if [[ -d "$target" ]]; then
        (cd "$target" && pwd -P)
    else
        local dir base
        dir="$(dirname "$target")"
        base="$(basename "$target")"
        (cd "$dir" && printf '%s/%s\n' "$(pwd -P)" "$base")
    fi
}

########################################
# Dependency checks
########################################

if ! command -v newstack >/dev/null 2>&1; then
    echo "ERROR: IMOD 'newstack' was not found in PATH." >&2
    exit 1
fi

########################################
# Read user inputs
########################################

read -r -p "Enter parent directory containing tilt-series folders: " parent_dir
parent_dir="$(trim "${parent_dir}")"

if [[ -z "${parent_dir}" ]]; then
    echo "ERROR: Parent directory is required." >&2
    exit 1
fi

if [[ ! -d "${parent_dir}" ]]; then
    echo "ERROR: Directory not found: ${parent_dir}" >&2
    exit 1
fi

parent_dir="$(abs_path "${parent_dir}")"

read -r -p "Enter output root directory [default: ${parent_dir}/processing]: " proc_dir
proc_dir="$(trim "${proc_dir}")"
if [[ -z "${proc_dir}" ]]; then
    proc_dir="${parent_dir}/processing"
fi
mkdir -p "${proc_dir}"
```

```
proc_dir="$(abs_path "${proc_dir}")"

read -r -p "Enter motion subdirectory name [default: motion]: " motion_subdir
motion_subdir="$(trim "${motion_subdir}")"
if [[ -z "${motion_subdir}" ]]; then
   motion_subdir="motion"
fi

read -r -p "Enter motion-corrected filename pattern [default: *_*.mc.mrc]: " mc_pattern
mc_pattern="$(trim "${mc_pattern}")"
if [[ -z "${mc_pattern}" ]]; then
   mc_pattern="*_*.mc.mrc"
fi

read -r -p "Enter sorted filelist name [default: sorted_filelist.txt]: " sorted_filelist_name
sorted_filelist_name="$(trim "${sorted_filelist_name}")"
if [[ -z "${sorted_filelist_name}" ]]; then
   sorted_filelist_name="sorted_filelist.txt"
fi

read -r -p "Enter output stack name [default: full_stack.mrc]: " stack_name
stack_name="$(trim "${stack_name}")"
if [[ -z "${stack_name}" ]]; then
   stack_name="full_stack.mrc"
fi

echo
echo "Parent directory : ${parent_dir}"
echo "Output root      : ${proc_dir}"
echo "Motion subdir    : ${motion_subdir}"
echo "MC file pattern  : ${mc_pattern}"
echo "Filelist name    : ${sorted_filelist_name}"
echo "Stack name       : ${stack_name}"
echo

########################################
# Main loop over immediate subdirectories
########################################

found_any=0
processed_any=0

for ts_dir in "${parent_dir}"/*; do
   [[ -e "${ts_dir}" ]] || continue
```

```
[[ -d "${ts_dir}" ]] || continue

# Skip the processing directory itself if it lives under parent_dir
if [[ "$(abs_path "${ts_dir}")" == "${proc_dir}" ]]; then
   continue
fi

found_any=1
ts_name="$(basename "${ts_dir}")"
ts_proc_dir="${proc_dir}/${ts_name}"
mkdir -p "${ts_proc_dir}"

motion_dir="${ts_dir}/${motion_subdir}"

echo "Processing: ${ts_name}"

if [[ ! -d "${motion_dir}" ]]; then
   echo "  WARNING: Motion directory not found: ${motion_dir}"
   echo "  Skipping."
   echo
   continue
fi

# Build the sorted file list by extracting the final underscore-delimited
# field before ".mc.mrc" as the tilt angle, then sorting numerically.
find "${motion_dir}" -maxdepth 1 -type f -name "${mc_pattern}" | \
   awk -F'_' '
      {
         angle = $NF
         sub(/\.mc\.mrc$/, "", angle)
         print angle "\t" $0
      }
   ' | \
   sort -g -k1,1 | \
   cut -f2- > "${ts_proc_dir}/${sorted_filelist_name}"

if [[ ! -s "${ts_proc_dir}/${sorted_filelist_name}" ]]; then
   echo "  WARNING: No files matched pattern ${mc_pattern}"
   echo "  Skipping."
   echo
   continue
fi

n_files="$(wc -l < "${ts_proc_dir}/${sorted_filelist_name}" | tr -d ' ')"
```

```
    echo "  Found ${n_files} tilt images"
    echo "  File list written: ${ts_proc_dir}/${sorted_filelist_name}"

    # Read the file list into a Bash array.
    # This lets us pass each file path directly to newstack.
    mapfile -t mc_files < "${ts_proc_dir}/${sorted_filelist_name}"

    if [[ "${#mc_files[@]}" -eq 0 ]]; then
        echo "  ERROR: File list was written but no entries were loaded." >&2
        echo
        continue
    fi

    echo "  Building stack with positional inputs..."

    # Remove any stale output before rebuilding
    rm -f "${ts_proc_dir}/${stack_name}"

    # Build the stack using positional input files followed by the output path.
    if newstack "${mc_files[@]}" "${ts_proc_dir}/${stack_name}"; then
        echo "  Stack created : ${ts_proc_dir}/${stack_name}"
        processed_any=1
    else
        echo "  ERROR: Stacking failed for ${ts_name}" >&2
        echo "  File list     : ${ts_proc_dir}/${sorted_filelist_name}" >&2
        echo "  Intended stack: ${ts_proc_dir}/${stack_name}" >&2
    fi

    echo
done

########################################
# Final report
########################################

if [[ "${found_any}" -eq 0 ]]; then
    echo "ERROR: No subdirectories were found under: ${parent_dir}" >&2
    exit 1
fi

if [[ "${processed_any}" -eq 0 ]]; then
    echo "ERROR: No tilt series were successfully stacked." >&2
    exit 1
fi
```

```
echo "Done."
echo "Stacks were written under: ${proc_dir}"
```
