## Supplementary material for "A robust workflow for 3D imaging of human mitochondria using cryo-electron tomography": Code 2

**Code 2. Scripts for removing bad tilts and writing new tilt angle file**

```
#!/usr/bin/env bash
#
# clean_tilt_stack_generic.sh
#
# Purpose:
#   Remove unwanted tilt images from an MRC tilt-series stack and generate
#   a matching tilt-angle list for the retained images.
#
# This version is path-independent:
#   - no hard-coded filenames
#   - no dependence on current working directory
#   - user provides both the input stack path and the sorted filelist path
#
# Inputs:
#   1. An input MRC stack
#   2. A sorted filelist used to build that stack
#
# Outputs:
#   <input_stem>_cleaned.mrc
#   <input_stem>_cleaned_TLT.txt
#
# Notes:
#   - The user enters frame numbers to remove using 1-based indexing.
#   - IMOD newstack expects excluded sections in 0-based indexing.
#   - If the user presses Enter without specifying any frames, the script
#     keeps all frames unchanged and writes the full TLT list.
#

set -euo pipefail
IFS=$'\n\t'

echo "=== Clean Tilt Stack and Export TLT ==="
echo

########################################
# Helper functions
########################################

# Trim leading/trailing whitespace from a string
trim() {
  local s="$1"
  s="${s#"${s%%[![:space:]]*}"}"
```

```
    s="${s%"${s##*[![:space:]]}"}"
    printf '%s' "$s"
}

# Convert an existing file or directory to an absolute physical path
abs_path() {
    local target="$1"
    if [[ -d "$target" ]]; then
        (cd "$target" && pwd -P)
    else
        local dir base
        dir="$(dirname "$target")"
        base="$(basename "$target")"
        (cd "$dir" && printf '%s/%s\n' "$(pwd -P)" "$base")
    fi
}

########################################
# Dependency checks
########################################

# Confirm that IMOD newstack is available
if ! command -v newstack >/dev/null 2>&1; then
    echo "ERROR: IMOD 'newstack' was not found in PATH." >&2
    exit 1
fi

########################################
# Read input stack path
########################################

# Ask the user for the input stack path
read -r -p "Enter input stack path (e.g., /path/to/full_stack.mrc): " in_mrc
in_mrc="$(trim "${in_mrc}")"

if [[ -z "${in_mrc}" ]]; then
    echo "ERROR: Input stack path is required." >&2
    exit 1
fi

if [[ ! -f "${in_mrc}" ]]; then
    echo "ERROR: Input stack not found: ${in_mrc}" >&2
    exit 1
fi
```

```
in_mrc="$(abs_path "${in_mrc}")"

########################################
# Read sorted filelist path
########################################

# Suggest a default sorted filelist in the same directory as the input stack
default_sorted_filelist="$(dirname "${in_mrc}")/sorted_filelist.txt"

read -r -p "Enter sorted filelist path [default: ${default_sorted_filelist}]: " sorted_filelist
sorted_filelist="$(trim "${sorted_filelist}")"
if [[ -z "${sorted_filelist}" ]]; then
   sorted_filelist="${default_sorted_filelist}"
fi

if [[ ! -f "${sorted_filelist}" ]]; then
   echo "ERROR: Sorted filelist not found: ${sorted_filelist}" >&2
   exit 1
fi

sorted_filelist="$(abs_path "${sorted_filelist}")"

########################################
# Define output filenames
########################################

# Build output names by appending "_cleaned" before the extension
base="${in_mrc%.*}"
ext="${in_mrc##*.}"

if [[ "${base}" == "${in_mrc}" ]]; then
   out_mrc="${in_mrc}_cleaned"
   out_stem="${out_mrc}"
else
   out_mrc="${base}_cleaned.${ext}"
   out_stem="${base}_cleaned"
fi

# Write the retained tilt angles to a matching TLT text file
out_tlt="${out_stem}_TLT.txt"

echo
echo "Input stack      : ${in_mrc}"
```

```
echo "Sorted filelist : ${sorted_filelist}"
echo "Output stack    : ${out_mrc}"
echo "Output TLT      : ${out_tlt}"
echo

########################################
# Extract all tilt angles from filenames
########################################

# Create a temporary file containing the full ordered angle list.
# This assumes each line of the sorted filelist ends with a filename whose
# final underscore-delimited field is the tilt angle before ".mc.mrc".
tmp_all_angles="$(mktemp)"

awk -F'_' '
   {
      angle = $NF
      sub(/\.mc\.mrc$/, "", angle)
      print angle
   }
' "${sorted_filelist}" > "${tmp_all_angles}"

# Count the number of frames represented in the filelist
total="$(wc -l < "${tmp_all_angles}" | tr -d ' ')"

if [[ "${total}" -eq 0 ]]; then
   echo "ERROR: Sorted filelist contains no entries: ${sorted_filelist}" >&2
   rm -f "${tmp_all_angles}"
   exit 1
fi

echo "Total frames detected: ${total}"
echo

########################################
# Read frames to exclude
########################################

# Ask the user which frames to remove.
# Accepted input examples:
#   3
#   11-17
#   3,13-15,30-34
read -r -p "Enter frames to remove (1-based; press Enter to keep all): " frames
```

```
frames="${frames// /}"

########################################
# Keep all frames if nothing was entered
########################################

# If the user enters nothing, preserve the full input stack and full angle list
if [[ -z "${frames}" ]]; then
   echo
   echo "No frames selected for removal."
   cp "${in_mrc}" "${out_mrc}"
   cp "${tmp_all_angles}" "${out_tlt}"
   rm -f "${tmp_all_angles}"

   echo "Done."
   echo "Retained all ${total} tilt angles:"
   cat "${out_tlt}"
   exit 0
fi

########################################
# Convert 1-based frame list to 0-based
# section list for newstack
########################################

# Convert the user's 1-based frame specification into the 0-based format
# expected by IMOD newstack -exclude
frames_0based="$(
echo "${frames}" | awk -F',' '
{
   for (i = 1; i <= NF; i++) {
      if ($i ~ /^[0-9]+-[0-9]+$/) {
         split($i, a, "-")
         if (a[1] < 1 || a[2] < 1 || a[2] < a[1]) {
            exit 1
         }
         printf "%d-%d", a[1] - 1, a[2] - 1
      }
      else if ($i ~ /^[0-9]+$/) {
         if ($i < 1) {
            exit 1
         }
         printf "%d", $i - 1
      }
```

```
        else {
            exit 2
        }

        if (i < NF) {
            printf ","
        }
    }
}
'
)" || {
    echo "ERROR: Could not parse frame specification: ${frames}" >&2
    rm -f "${tmp_all_angles}"
    exit 1
}

echo
echo "Removing frames (0-based for newstack): ${frames_0based}"

########################################
# Create cleaned stack
########################################

# Exclude the requested sections from the input stack
newstack -input "${in_mrc}" -output "${out_mrc}" -exclude "${frames_0based}"

########################################
# Build retained tilt-angle list
########################################

# Build a sed deletion command using the original 1-based frame numbers,
# so the retained TLT file stays synchronized with the cleaned stack.
sed_cmd=""
IFS=',' read -r -a parts <<< "${frames}"

for p in "${parts[@]}"; do
    if [[ "${p}" =~ ^[0-9]+-[0-9]+$ ]]; then
        a="${p%-*}"
        b="${p#*-}"

        if (( a < 1 || b < a )); then
            echo "ERROR: Invalid frame range: ${p}" >&2
            rm -f "${tmp_all_angles}"
            exit 1
```

```
        fi

        for ((k=a; k<=b; k++)); do
            sed_cmd+="${k}d;"
        done
    elif [[ "${p}" =~ ^[0-9]+$ ]]; then
        if (( p < 1 )); then
            echo "ERROR: Invalid frame number: ${p}" >&2
            rm -f "${tmp_all_angles}"
            exit 1
        fi

        sed_cmd+="${p}d;"
    else
        echo "ERROR: Could not parse frame token: '${p}'" >&2
        rm -f "${tmp_all_angles}"
        exit 1
    fi
done

# Write the retained tilt angles after removing the excluded frames
sed "${sed_cmd}" "${tmp_all_angles}" > "${out_tlt}"

# Clean up the temporary file
rm -f "${tmp_all_angles}"

########################################
# Final report
########################################

kept="$(wc -l < "${out_tlt}" | tr -d ' ')"

echo
echo "Done."
echo "Retained ${kept} tilt angles:"
cat "${out_tlt}"
```
